# CMTM6 mediates cisplatin resistance in OSCC by regulating AKT/c-MYC driven ribosome biogenesis

**DOI:** 10.1101/2022.04.08.487634

**Authors:** Pallavi Mohapatra, Sibasish Mohanty, Shamima Azma Ansari, Omprakash Shriwas, Arup Ghosh, Rachna Rath, Saroj Kumar Das Majumdar, Rajeeb K Swain, Sunil K Raghav, Rupesh Dash

## Abstract

CMTM6, a type 3 transmembrane protein, is known to stabilize the expression of programmed cell death ligand 1 (PD-L1) and hence facilitates the immune evasion of tumor cells. Recently, we demonstrated that CMTM6 is a major driver of cisplatin resistance in oral squamous cell carcinomas (OSCC). However, the detailed mechanism how CMTM6 rewires cisplatin resistance in OSCC is yet to be explored. RNA sequencing analysis of cisplatin resistant OSCC lines stably expressing NtShRNA and CMTM6 ShRNA revealed that CMTM6 might be a potential regulator of ribosome biogenesis network. Knocking down CMTM6 significantly inhibited transcription of 47S precursor rRNA and hindered the nucleolar structure, indicating reduced ribosome biogenesis. When CMTM6 was ectopically over expressed in CMTM6KD cells, almost all ribosomal machinery components were rescued. Mechanistically, CMTM6 induced the expression of C-Myc, which promotes RNA polymerase I mediated rDNA transcription. In addition to this, CMTM6 also found to regulate the AKT–mTORC1-dependent ribosome biogenesis and protein synthesis in cisplatin resistant lines. The nude mice and zebrafish xenograft experiments indicate that blocking ribosome synthesis either by genetic inhibitor (CMTM6KD) or by pharmacological inhibitor (CX-5461), significantly restores cisplatin medicated cell death in chemoresistant OSCC. Overall, our study suggests that CMTM6 is a major regulator of ribosome biogenesis network and targeting ribosome biogenesis network is a viable target to overcome chemoresistance in OSCC. The novel combination of CX-5461 and cisplatin deserves further clinical investigation in advanced OSCC.

## Introduction

Squamous cell carcinomas accounts for 90% of head and neck cancers worldwide (1). In India, OSCC is the most prevalent cancer among male with almost 80,000 new cases reported each year (2). Unfortunately, 70% of the OSCC cases are diagnosed at advanced stage without having any history of preclinical lesion. The most common treatment regimens are surgery followed by concurrent chemo radiotherapy. Neoadjuvant chemotherapy is frequently prescribed for surgically unresectable OSCC tumors (3). In spite of having these treatment modalities, the 5 year survival rate of advanced tongue squamous cell carcinomas is less than 5 years. Chemoresistance is one of the important factor for treatment failure in OSCC. Cisplatin, 5FU and docetaxel (TPF) are most common chemotherapy regimen used for OSCC. Though TPF regimen shows initial positive result, the tumour progressively acquires drug resistance and hence patients experience continued tumor growth and metastatic disease (4). The molecular mechanism behind the chemoresistance in OSCC is poorly understood. To explore the causative factors responsible for cisplatin resistance, we earlier performed global proteomic profiling of human OSCC lines presenting with sensitive, early and late cisplatin resistance patterns. Our study suggests that CMTM6 modulates cisplatin resistance by activating Wnt signalling and upregulating proto-oncogene C-Myc (5). However, the exact molecular mechanism by which CMTM6 regulates cisplatin resistance is still to be explored.

Enhanced rate of ribosome biogenesis is frequently observed during malignant transformation and hence imbalance in ribosome function and synthesis is common event during the process of carcinogenesis (6). The rDNA transcription occurs in nucleolus and hence nucleolus serves as the site for ribosome biogenesis inside the cells. The enhanced size of nucleolus or hypertrophied nucleoli in tumor tissues (as compared to non-malignant adjacent tissues) is a hallmark of tumor malignancy (7). The ultrastructure of nucleolus revealed that the rRNA genes are located in fibrillar components of nucleolus along with all the necessary factors required for rDNA transcription (8). The transcribed rRNA are confound in dense fibrillar component. Hence, the structural organization of fibrillar components represent the status of ribosome biogenesis in a cell (9). The rDNA is transcribed by RNA Pol I to produce 47S precursor rRNA. Next, the 47S rRNA undergoes various modifications to eliminate the external (5′- and 3′-ETS) and internal (ITS 1and 2) transcribed spacers to yield the mature rRNA i.e., 28S,18S and 5.8S rRNA. The 5’ETS plays important role in initiation of transcription of rDNA (10). Three different important factors are essential for RNA Pol I to perform rDNA transcription, those are termed transcription initiation factor I (TIF-I), selectivity factor 1 (SL1), and upstream binding factor (UBF). Similarly, RNA Pol III transcribes 5S rRNA in nucleoplasm and ultimately transported to nucleolus (11). The rRNA are assembled with 79 different types of ribosomal proteins (RPs) to generate mature ribosomal subunits. The 18S rRNA assembled with 33 different small ribosomal proteins (RPSs) to produce 40S subunit, whereas 60S rRNA subunit consists of one each of the 28S,5.8S, and 5S rRNAs, along with 47 large ribosomal proteins (RPLs) (11). Finally, both 40S and 60S rRNA subunits translocate to cytoplasm to form 80S ribosomal particles. Since, enhanced ribosome biogenesis often observed in malignant cells, efforts have been made to discover small molecules those block rDNA transcription and hence developing a strategy to target cancer cells. The most potent inhibitor of rRNA synthesis discovered till date is CX-5461, which is a benzothiazole compound that selectively inhibits activity RNA Pol I. CX5461 supressed the binding of transcription factors of RNA Pol I to rDNA promoters, particularly CX5461 significantly blocks the binding of SL1 and decreases the Pol I activity by 40-60% (12). Hence, CX-5461 blocks the growth and proliferation of leukaemia, lymphoma, colorectal and pancreatic carcinomas (13,14). A phase I clinical trial has been successfully conducted for haematological malignancies and currently Phase I/II clinical trial is undergoing for solid tumors.

Our interest in studying ribosome biogenesis came from the finding that ribosome biogenesis network is deregulated when we analysed the transcriptome data of OSCC chemoresistant lines stably expressing NtShRNA and CMTM6ShRNA (KD). Earlier, other groups have suggested that CMTM6 is known to stabilize the expression of programmed cell death ligand 1 (PD-L1) and hence facilitates the immune evasion of tumor cells (15,16). We have also recently demonstrated that CMTM6 is a major driver of cisplatin resistance in OSCC (5). In this study, we wanted to explore I) If CMTM6 is a potential regulator of ribosome biogenesis network in OSCC chemoresistant cells II) if targeting ribosome biogenesis network is a viable target to overcome cisplatin resistance in OSCC.

## Result

### Transcriptome analysis revealed CMTM6 as a potential regulator of ribosome biogenesis

Recently, employing global proteomic profiling of cisplatin sensitive, early and late resistant OSCC cell lines, we revealed CMTM6 as a major driver of cisplatin resistance in OSCC. In the same study, our transcriptome analysis of control and CMTM6 depleted cisplatin resistant OSCC lines in untreated and cisplatin treatment condition (5) showed “Wnt Signaling” as one of the top enriched pathways based on the number of genes involved in the process (5). When we arranged the enriched GO list based on p-value significance (p<0.05, p adjusted <0.05) the top significantly depicted pathway was “Ribosome Biogenesis” and “ribonucleoprotein complex biogenesis” in the CMTM6 depleted as compared to control vector transfected cells (Figure 1 A-D). The datasets generated in the study are submitted and available in EBI Array Express portal with the accession id E-MTAB-9424. To confirm the clinical relevance of our finding from RNA-sequencing data, we performed association analysis between CMTM6 and genes found in figure 1D by online databases of cancer genome atlas HNSCC cohort using GEPIA. The association analysis of CMTM6 mRNA levels with 35 randomly selected ribosome biosynthesis related genes found in figure 1D from the HNSCC cohort showed a positive correlation (r > 0.2) (Supplemental Figure 1), which suggests that CMTM6 potentially regulates ribosome biosynthesis.

**Figure 1:**
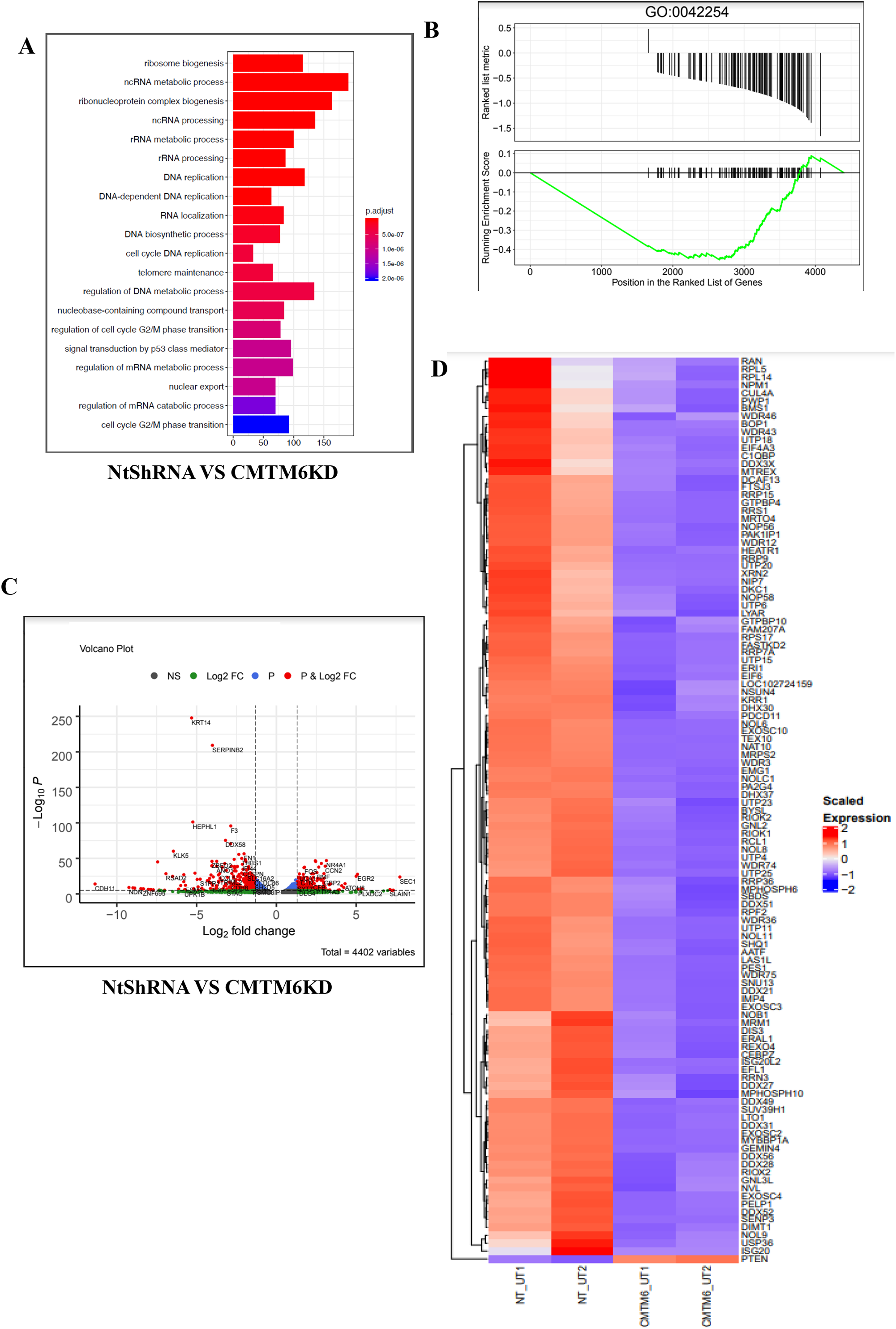
Transcriptome analysis of untreated control and CMTM6KD cisplatin resistant OSCC lines revealed CMTM6 as a potential regulator of ribosome biogenesis. A) Top twenty gene ontology terms (GO molecular function) enriched in differentially expressed genes of NtShRNA versus CMTM6 knockdown (CMTM6KD) cisplatin resistant OSCC line (H357CisR). The GO terms are arranged based on their adjusted p-values B) Gene set enrichment analysis (GSEA) plot of NtShRNA versus CMTM6KD cells for genes involved in ribosome biogenesis (GO:0042254), C) Volcano plot of ribosome biogenesis pathway genes in untreated control versus CMTM6 knockdown H357 cisplatin resistant cancer cell line with select. Few of the ribosome biogenesis genes are marked in the plot, D) Heatmap representing the expression pattens of genes involved in ribosome biogenesis in untreated control vector and CMTM6 knockdown RNA-seq (n=2).

### CMTM6 regulates ribosome biogenesis in cisplatin resistant OSCC

To validate the finding from transcriptomic analysis, qPCR was performed to measure the expression of 5’ETS of 47S precursor rRNA in cisplatin resistant OSCC lines stably expressing NtShRNA and CMTM6ShRNA#2 (KD). The data suggest that there is a significant decrease of newly transcribed 47S precursor rRNA in CMTM6 KD cells, which was efficiently rescued with ectopic overexpression of CMTM6 (Figure 2A, B). Here it is important to mention that we earlier generated cisplatin resistant OSCC lines (CisR) with prolonged treatment of cisplatin to OSCC lines (17) and to knock down CMTM6, we used shRNA sequence those targeting 5′UTR (CMTM6 shRNA#2) of CMTM6 mRNA. This clone was generated in our previous study (5) and here onwards we will term it as CMTM6KD. Next, we found that the relative expression of mature rRNAs (28S,18S and 5.8S) are significantly lower in CMTM6KD cells as compared to NtShRNA in chemoresistant OSCC lines (Figure 2C-D). We also achieved similar observation with another CMTM6ShRNA#1 which targets the coding sequence of CMTM6 mRNA (Supplementary Figure 2). Here onwards, for rest of the experiments we have used CMTM6 shRNA#2 (CMTM6KD). Similarly, in Patient-derived primary tumor cells (PDC1) not responding to taxol and platinum therapy (TP), CMTM6 knock down results in decreased abundance of 5’ETS and mature rRNAs (Figure 2B and 2E). Analysis of clinical samples revealed that, abundance of 47S 5’ETS and mature rRNA are significantly higher in tumors of chemotherapy (TPF) non-responder OSCC patients as compared to chemotherapy responders, which is positively correlated with CMTM6 expression (Figure 2F-M). Next, we monitored nucleolar morphology by immunostaining of Fibrillarin protein. The data suggest that knocking down CMTM6 results in dispersal of Fibrillarin and the loss of multiple nucleoli, which is rescued by ectopic overexpression of CMTM6 (Figure 2N). Here we treated CX-5461 as a positive control, which inhibits rDNA transcription. In CX5461 treated group, we observed loss of nucleolus indicating reduced rRNA synthesis (Figure 2N).

**Figure 2:**
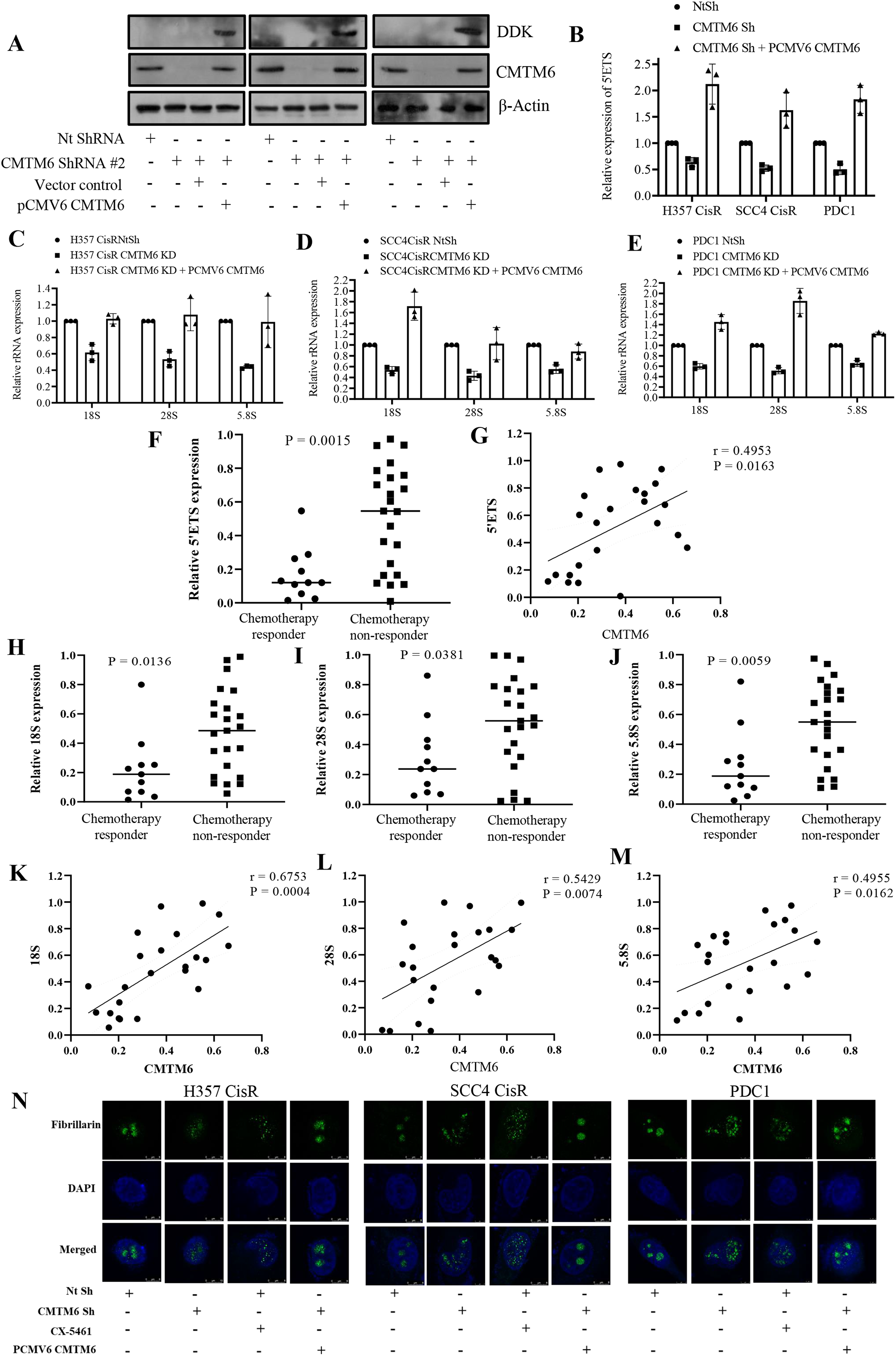
CMTM6 acts as a regulator of ribosome biogenesis in cisplatin resistant OSCC: **A)** Cisplatin resistant OSCC lines were stably transfected with NTShRNA and CMTM6ShRNA. ShRNA#2 targets 5’UTR of CMTM6 mRNA. For ectopic overexpression, pCMV6-Entry-CMTM6 (MYC-DDK tagged) and control vector were transiently transfected to indicated CMTMKD (ShRNA#2) cells and immunoblotting (n=3) was performed with indicated antibody of indicated genes. Efficient overexpression was evident from CMTM6 and DDK expression. **B)** CMTM6 was transiently over expressed in chemoresistant cells stably expressing CMTM6ShRNA#2 and relative 5’ETS and rRNA (fold change) expression analyzed by qRT PCR in indicted cells (mean ±SEM, n=3), *: P < 0.05 by two way ANOVA. **C-E)** CMTM6 was transiently over expressed in indicated chemoresistant cells stably expressing CMTM6ShRNA#2 and relative expression of 18S, 28S and 5.8S (fold change) were analyzed by qRT PCR in indicted cells (mean ±SEM, n=3). **F-G)** Relative 5’ETS expression was analyzed by qRT PCR in different chemotherapy-non-responder OSCC tumors as compared to chemotherapy responder tumors (Median, n=11 for chemotherapy responder and n=23 for chemotherapy-non-responder). *: P < 0.05 by Student’s t test and correlation analysis was done with CMTM6 expression. **H-J)** Relative 18S,28S and 5.8S rRNA expression were analyzed by qRT PCR in different chemotherapy-non-responder OSCC tumors as compared to chemotherapy responder tumors (Median, n=11 for chemotherapy responder and n=23 for chemotherapy-non-responder). *: P < 0.05 by Student’s t test and **K-M)** correlation analysis was done with CMTM6 expression. **N)** Chemoresistant cells stably expressing NTShRNA or CMTM6ShRNA#2 and pCMV6-Entry-CMTM6 were subjected to immunostaining and confocal microscopy with indicated antibodies depicting nucleolar shape.

### CMTM6 regulates ribosome biogenesis through c-Myc and AKT axis

In our previous report, we have shown that CMTM6 regulates the Wnt signalling and its target genes including proto-oncogene c-Myc (5). Interestingly, c-Myc is known to tightly regulate the Pol I activity and thereby modulates rDNA transcription. Hence, we hypothesize that CMTM6 regulates ribosome biogenesis by inducing c-Myc expression. In CMTM6KD cells, we found reduced expression of c-Myc, which was efficiently rescued by ectopic expression of CMTM6 (Figure 3A). However, when we ectopically over expressed c-Myc in CMTM6KD cells we did not find any alteration of CMTM6 expression (Figure 3B). These observations indicate that CMTM6 is upstream to c-Myc. Next, we scored the abundance of 5’ETS in 47S rRNA and mature rRNA in CMTM6KD cells with ectopic overexpression of c-Myc. The data suggest that the rRNA synthesis is reduced in CMTM6KD cells, which is rescued with ectopic overexpression of c-Myc (Figure 3C-E). In clinical samples, we found the expression of c-Myc to be elevated in the tumor tissues of chemotherapy non-responders as compared to responders, where a clear correlation could be found with CMTM6 (Figure 3F, G). Similarly, ectopic overexpression of c-Myc efficiently rescued the nucleolar morphology in CMTM6KD cells (Figure 3H). In addition to c-Myc, we also earlier reported that CMTM6 interacts with Enolase-1 and modulates the activation of AKT (5). Since the PI3K–AKT–mTOR axis conjoin with MYC to control ribosome biogenesis, we explored the possible role of AKT in CMTM6 mediated ribosome biogenesis. First, in CMTM6KD cells there is a reduction of p-AKT (Ser473), which can be efficiently rescued with overexpression of CMTM6 (Figure 4A). Similarly, when myrAKT (constitutively active AKT) is ectopically overexpressed in CMTM6KD cells, we found rescue of c-MYC expression (Figure 4B). Again, inhibition of AKT by AKTi in CMTM6 overexpressing cells reduced c-MYC expression (Supplementary figure 3). Our data suggest that myr-AKT expression in CMTM6-KD cells efficiently rescued the level of 5’ETS and mature rRNA (Figure 4C-E). Again, with ectopic overexpression of myr-AKT, we observed partial rescue of nucleolar morphology in CMTM6KD cells (Figure 4F).

**Figure 3:**
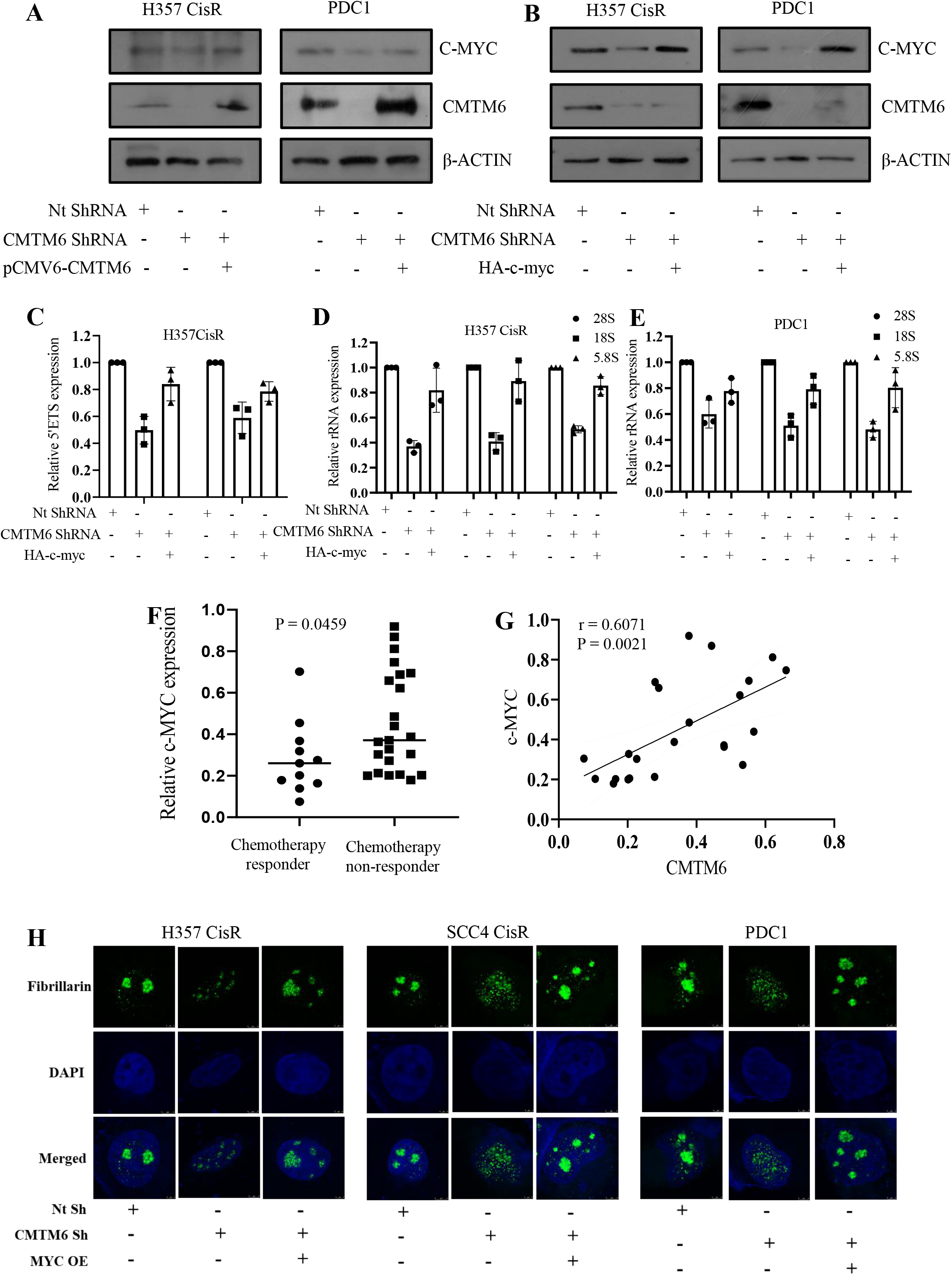
c-MYC regulates ribosome biogenesis in CMTM6 dependent manner: **A)** CMTM6 was overexpressed in chemoresistant cells stably expressing NTShRNA or CMTM6ShRNA#2 and immunoblotting was performed with indicated antibodies. **B)** c-MYC was overexpressed in chemoresistant cells stably expressing NTShRNA or CMTM6ShRNA#2 and immunoblotting was performed with indicated antibodies. **C-E)** c-MYC was transiently over expressed in chemoresistant cells stably expressing CMTM6ShRNA#2 and relative 5’ETS and mature rRNA (fold change) expression analyzed by qRT PCR in indicted cells (mean ±SEM, n=3), *: P < 0.05 by two way ANOVA. **F)** Relative c-MYC expression was analyzed by qRT PCR in different chemotherapy-non-responder OSCC tumors as compared to chemotherapy responder tumors (Median, n=11 for chemotherapy responder and n=23 for chemotherapy-non-responder). *: P < 0.05 by Student’s t test **G)** correlation analysis was done with CMTM6 expression. **H)** Chemoresistant cells stably expressing NTShRNA or CMTM6ShRNA#2 were subjected to immunostaining and confocal microscopy with indicated antibodies depicting nucleolar shape. Here MYC was transiently overexpressed in CMTM6KD cells.

**Figure 4:**
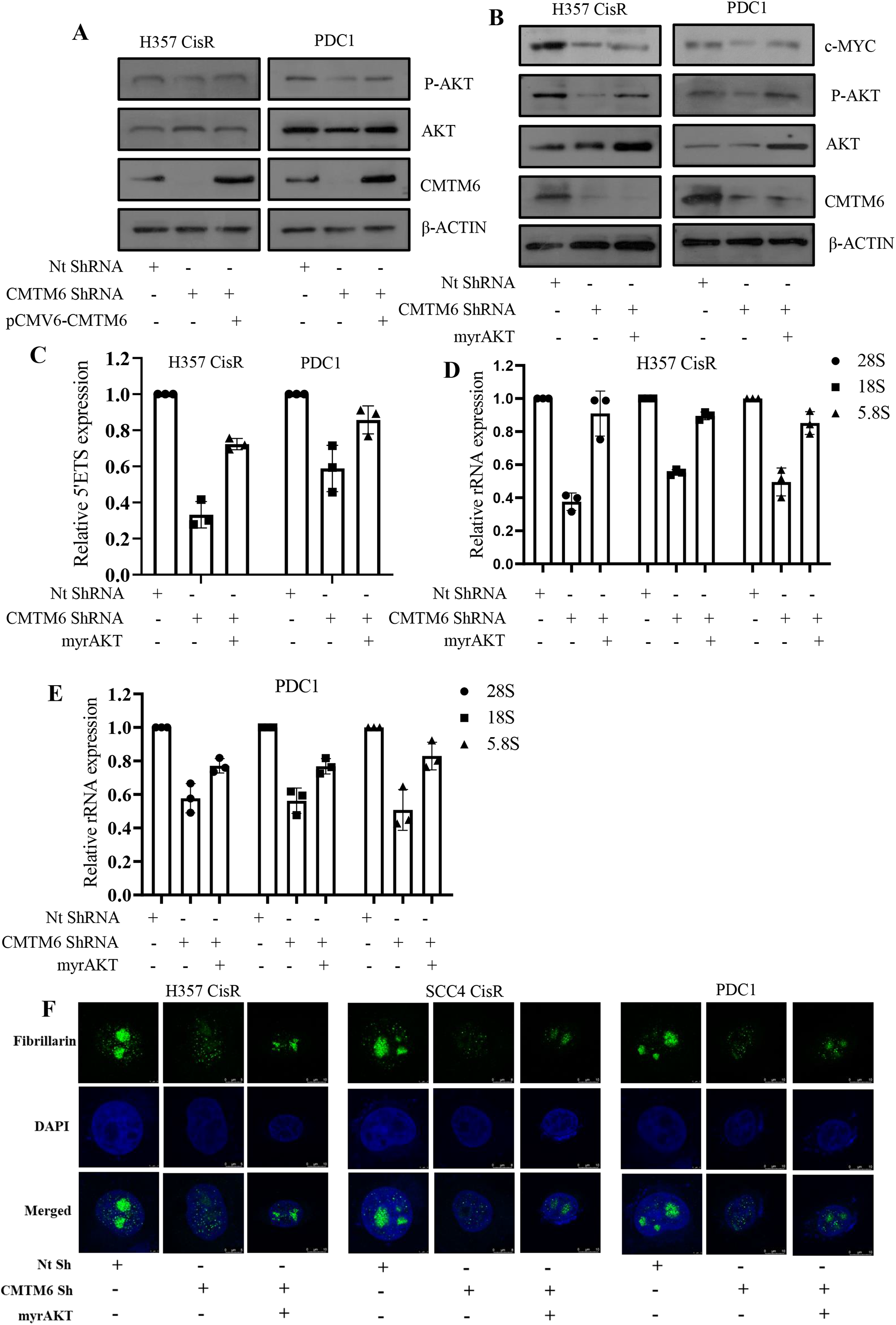
CMTM6 regulates AKT and thereby ribosome biogenesis: **A)** CMTM6 was overexpressed in chemoresistant cells stably expressing NTShRNA or CMTM6ShRNA#2 and immunoblotting was performed with indicated antibodies. **B)** myr-AKT was overexpressed in chemoresistant cells stably expressing NTShRNA or CMTM6ShRNA#2 and immunoblotting was performed with indicated antibodies. **C-E)** myr-AKT was transiently over expressed in chemoresistant cells stably expressing CMTM6ShRNA#2 and relative 5’ETS and rRNA (fold change) expression analyzed by qRT PCR in indicted cells (mean ±SEM, n=3), *: P < 0.05 by two way ANOVA. **F)** Chemoresistant cells stably expressing NTShRNA or CMTM6ShRNA#2 were subjected to immunostaining and confocal microscopy with indicated antibodies depicting nucleolar shape. Here myrAKT was transiently overexpressed in CMTM6KD cells.

### CMTM6 drives ribosome biogenesis via AKT/mTOR axis

The PI3K/AKT/mTOR pathway is activated by growth factors and hormones, which phosphorylates the ribosomal proteins S6 (RPS6) that in turn phosphorylates 4E-BP1. Phosphorylated 4EBP1 triggers its release from elongation factor eIF4E ensuring in enhanced cap-dependent translation. When we explored the AKT/mTOR axis in CMTM6KD cells, we found that in CMTM6KD cells there is a reduction in expression of p-mTOR, p-RPS6 and p-4E-BP1, which is significantly rescued by ectopic overexpression of CMTM6 (Figure 5A). Inhibition of AKT by AKTi resulted in reduced expression of p-mTOR, p-RPS6 and p-4E-BP1 in presence of CMTM6, whereas overexpression of AKT in CMTM6KD cell displayed a reversed effect, suggesting the AKT dependency of CMTM6 for the same (Figure 5B). Again, inhibition of mTOR activity by rapamycin resulted in reduced expression of p-RPS6 and p-4EBP1 in the CMTM6 or AKT overexpressed cells demonstrating the CMTM6-AKT-mTOR axis (Figure 5C). Overall, these studies suggest that CMTM6 regulates ribosome biogenesis and protein synthesis network through c-Myc and AKT. Finally, to confirm the impact of CMTM6 in translation, that is denoted as the last step of ribosome biogenesis, we performed puromycin incorporation assay. Here, actively growing cisplatin resistance lines stably expressing NtShRNA and CMTM6ShRNA were incubated with a short pulse of puromycin (15 min, 10 μM). Puromycin (an analog of tyrosyl-tRNA) gets incorporated into nascent polypeptide chain. Hence, immunoblotting of lysates isolated from puromycin treated cells with anti-puromycin antibody provides an estimation of nascent translation rate. In harmony with our hypothesis, the translation was inhibited in CMTM6KD cells as compared to NtShRNA cells, which was rescued with ectopic overexpression of CMTM6 (Figure 5D).

**Figure 5:**
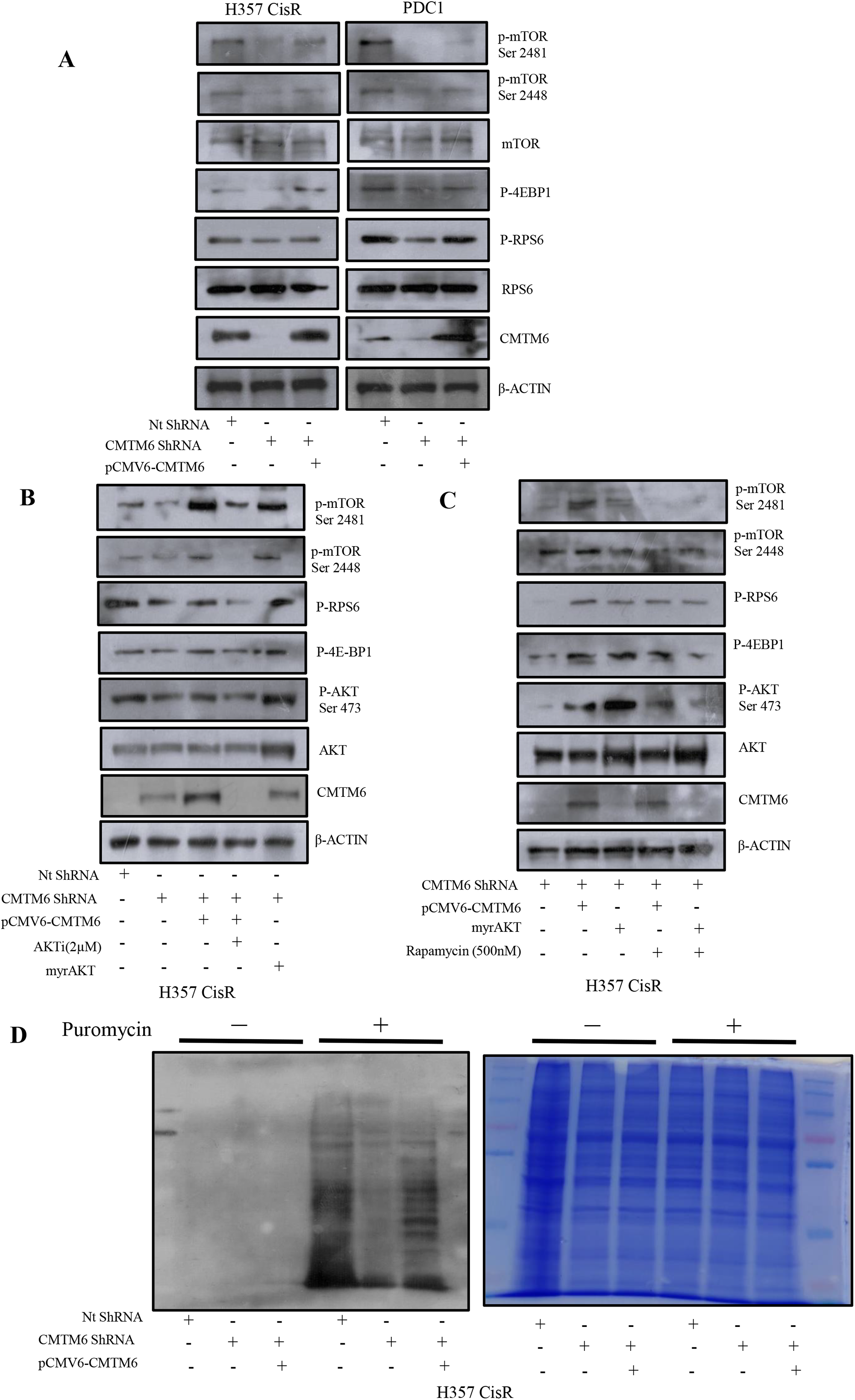
CMTM6 regulates AKT/mTOR axis and assembly of ribosome: **A)** CMTM6 was overexpressed in chemoresistant cells stably expressing CMTM6ShRNA#2 and immunoblotting was performed with indicated antibodies. **B)** Lysates were isolated from indicated AKTi treated cells and vehicle control and immunoblotting was performed with indicated antibodies. **C)** Lysates were isolated from indicated Rapamycin treated cells and vehicle control and immunoblotting was performed with indicated antibodies. **D)** Lysates were isolated from puromycin treated (15 mins pulse) cells and vehicle control. Left panel: Immunoblotting (n=3) was performed using indicated antibodies. Right panel: Coomassie brilliant blue stanning was performed and photograph was taken.

### Supressing ribosome biogenesis restores cisplatin induced cell death in chemoresistant OSCC

To understand the consequences of our finding that CMTM6 regulates ribosome biogenesis in the context of drug resistance, we checked if supressing ribosome biogenesis by either pharmacological inhibitor (CX-5461) or genetic inhibition (CMTM6KD) could reverse cisplatin resistance. First, we determined the IC50 value of CX-5461 in chemoresistant OSCC lines. The data suggest that the IC50 is approximately 7.5 μM for all three lines i.e. SCC4 CisR, H357CisR and PDC1 (Figure 6A). For combinatorial treatment, we considered non-toxic dose of CX-5461 i.e., 0.5 and 1 μM which is enough to inhibit rDNA transcription (Supplementary figure 4). The cell viability, cell death, colony forming assay and immunoblotting data revealed that non-toxic dose of CX-5461 efficiently restored cisplatin mediated cell death in cisplatin resistant OSCC (Figure 6 B-E). Similar result is generated with treatment of cisplatin in CMTM6KD cells (Figure 6B, C and E).

**Figure 6:**
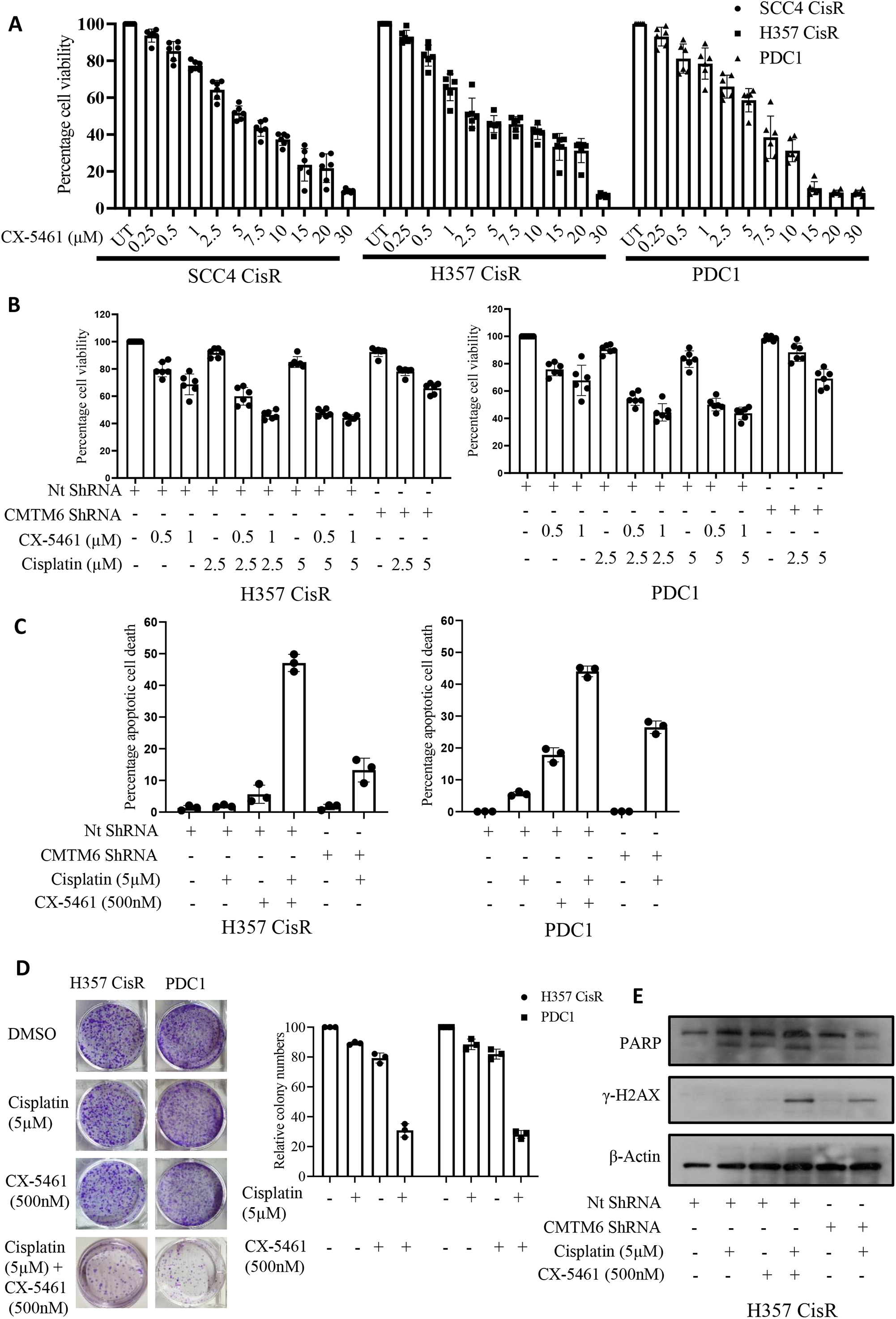
Treatment of POL-I inhibitor CX-5461 reverses cisplatin resistant phenotype: **A)** Indicated cells were treated with CX-5461 for 48h with indicated concentration and cell viability was determined by MTT assay (n=3), *: P < 0.05. **B)** Chemoresistant cells were treated with indicated concentration of Cisplatin and CX-5461 for 48h with indicated concentration and cell viability was determined by MTT assay (n=3), *: P < 0.05. **C)** Chemoresistant cells were treated with indicated concentration of cisplatin and CX-5461 for 48h and cell death was determined by annexin V/7AAD assay using flow cytometer. (Mean ±SEM, n=3), *: P < 0.05. **D)** Right panel: Indicated cells were treated with cisplatin and CX-5461 for 12 days for colony forming assay (n=3 and *: P < 0.05). Left panel: representation photographs of colony forming assay in each group. **E)** Indicated cells were treated with cisplatin and CX-5461 for 48h and immunoblotting (n=3) was performed with indicated antibodies.

### Inhibition of ribosome biogenesis significantly induced cisplatin-mediated apoptosis in chemo-resistant patient-derived xenograft

We checked combinatorial efficacy of cisplatin and CX-5461 in nude mice xenograft model using PDC1 lines. The *in vivo* data suggest that the combination of CX-5461 (20mg/kg) and cisplatin (3 mg/kg) significantly reduced the tumor burden in the flank of nude mice as compared to treatment with either of the single agents (7A-C). Analysing the expression of Ki-67 in paraffin embedded tumor tissues, profound reduction of Ki-67 in combination group as compared to single treated group was observed (Figure 7D). Similarly, we used Zebrafish (*Danio rerio*) [Tg(fli1:EGFP)] tumor xenograft model to further validate our findings. Equal number of the PDC1 cells in four different groups were stained with Dil (1,1’-Dioctadecyl-3,3,3’,3’-Tetramethylindocarbocyanine Perchlorate) and injected into perivitelline space of 48-hour post fertilized zebrafish embryos. After 3 days of injection, embryos were treated with vehicle control or cisplatin (500ng/ml) or CX-5461 (500nM) or cisplatin in combination with CX-5461. After 5 days of injection, tumors were documented using a fluorescence microscope. The tumor growth, as measured by fluorescent intensity of primary tumors, was found to be significantly reduced in the combination group as compared to either of single treatment groups (Figure 7 E-F). In zebrafish tumor xenograft model, we also found that when cisplatin (500ng/ml) is treated to CMTM6 KD cells, there is a reduction of the tumor growth (Figure 7G and Supplementary figure 5). Overall, our data suggest that targeting ribosome biogenesis could be a viable option to overcome chemoresistance in OSCC.

**Figure 7:**
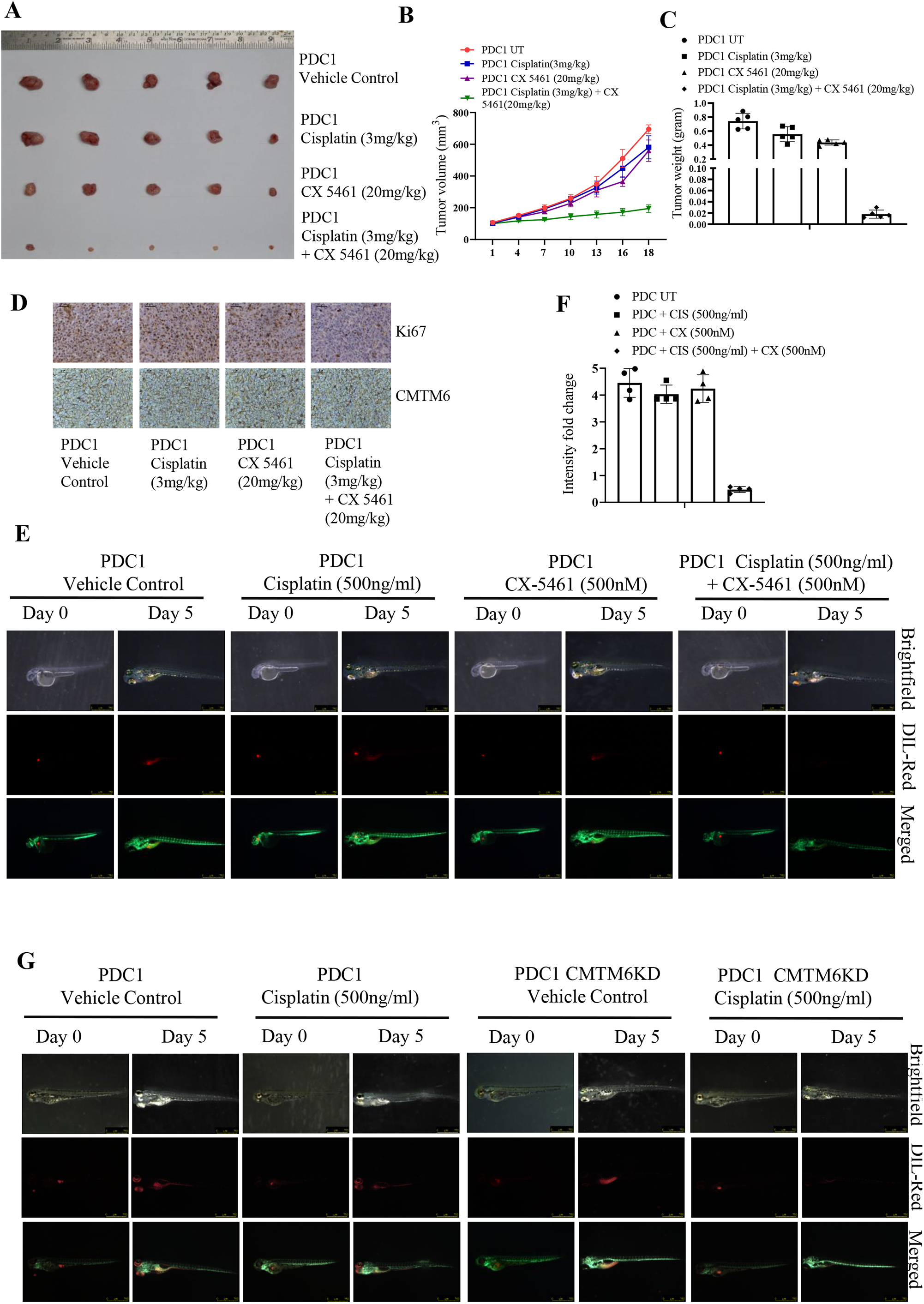
In-vivo validation of inhibition of ribosome biogenesis and cisplatin sensitivity: **A)** Patient derived cells (PDC1) established from tumor of chemo-nonresponder patient. PDC1 cells were implanted in right upper flank of athymic male nude mice, after which they were treated with cisplatin and CX-5461 at indicated concentration. At the end of the experiment mice were euthanized, tumors were isolated and photographed (n=5). **B)** Tumor growth was measured (mean ± SEM, n = 5), *: P < 0.05. **C)** Tumor weight measured at the end of the experiment (mean ± SEM, n = 5), *: P < 0.05. **D)** Tumors were isolated and paraffin-embedded sections were prepared to perform immunohistochemistry with indicated antibodies. **E)** Lateral view of fluorescent transgenic [Tg(fli1:EGFP)] zebrafish embryos at Day 0 and Day 5 injected with Dil-Red stained PDC1 cells with and without treatment of Cisplatin and CX-5461. The tumor growth and migration was assessed by an increase in fluorescence intensity on the 5th day compared to the day of injection. n=4. **F)** The quantitation of fluorescence intensity was performed using ImageJ software. **G)** Lateral view of fluorescent transgenic [Tg(fli1:EGFP)] zebrafish embryos at Day 0 and Day 5 injected with Dil-Red stained PDC1 control and CMTM6 KD cells with and without treatment of Cisplatin. The tumor growth was assessed by an increase in fluorescence intensity on the 5th day compared to the day of injection. n=4.

## Discussion

Myc is a master transcription factor which regulates the transcription of approximately 15% of the genome (18). It controls several biological processes and signalling pathways including many malignant phenotype like cell proliferation, cell death, angiogenesis, metastasis and metabolism. Ribosome biogenesis is one of the potent biological process controlled by Myc (19). It was observed that when Myc is ectopically overexpressed in mammalian cells, protein synthesis rate was dramatically increased along with cell proliferation and growth. Similarly, overexpression of diminutive (the drosophila Myc gene) in flies results in markedly enhanced nucleolar and cell size (20). B lymphocyte specific Myc overexpression in a transgenic mice resulted in increased protein synthesis and ribosomal biogenesis (21). Similarly, when Myc was ectopically overexpressed through an adenovirus in hepatocytes of mice, the nucleolar size was enhanced with marked up regulation of rDNA transcription (22). Interestingly, Myc modulates ribosome synthesis by regulating each of the steps of biogenesis i.e transcription, rRNA processing, assembly and translation. Myc regulates the rDNA transcription as it enhances the expression of RNA Pol I cofactors i.e. upstream binding transcription factor (UBF) and selectivity factor 1 (SL1). Myc interacts with transcription factor IIIB (TFIIIB) which is an essential cofactor for RNA Pol III to induce the transcription of 5S rRNA (23). Myc also regulates the activity of RNA Pol II and hence it tightly controls the transcription of several RPS and RPL (ribosomal proteins).

Overall, myc regulates the transcription of various proteins those play crucial role in ribosome biogenesis and translation initiation. Other than Myc, AKT/m-TOR pathway, the central regulator of translation, also regulates the rDNA transcription. AKT/m-TOR indirectly regulates TIF-1A, which is an important factor for RNA Pol I. Blocking m-TOR pathway increases the phosphorylation of TIF-1A at Ser199 and reduces the phosphorylation at Ser44 (24). TIF-1 knock out cells showed decreased ribosome biosynthesis machinery with reduced translation and decreased nucleolar size. Another study revealed that m-TOR also phosphorylates UBF, which is another important transcription factor for RNA Pol I (25). So it is broadly hypothesized that multipoint inhibition of both the important network of ribosome biogenesis (Myc and AKT-m-TOR) could be a potent approach to target cancer cells. Devlin et.al. have demonstrated recently that the combination of CX-5461 (inhibit rDNA transcription) and everolimus (inhibitor of m-TOR) synergistically induced apoptosis in Myc driven Tp53WT Eμ-Myc B-lymphoma cancer lines. *In vivo* experiments by transplanting Eμ-Myc in C57BL/6 mice suggest that length of the survival extension was double as compared to single treatment (26). In high-grade ovarian cancer PDX model involving chemosensitive and chemoresistant populations, CX-5641 (at 50 mg/kg) treatment showed variable anti-tumor activity i.e., out of five PDX, three did not show any response and two PDX showed response (27). These data suggests combining that CX-5461 and chemotherapeutic drug could be a viable approach to overcome chemoresistance.

Earlier, characterizing the cisplatin sensitive and resistant OSCC lines, we found that CMTM6 drives cisplatin resistance. Mechanistically, cisplatin binds to Enolase-1, which led to activation of Wnt/β-catenin signalling through AKT. As a consequence of this, several Wnt signalling target genes were regulated by CMTM6 including c-Myc. In this study, we have shown that CMTM6 regulates protein synthesis through a multipoint inhibition of ribosome biogenesis network i.e. CMTM6 regulates c-Myc as well as AKT/m-TOR pathway. Our study suggests that CMTM6 activates AKT and hence regulates c-Myc and m-TOR to modulate ribosome biogenesis and protein translation (Figure 8).

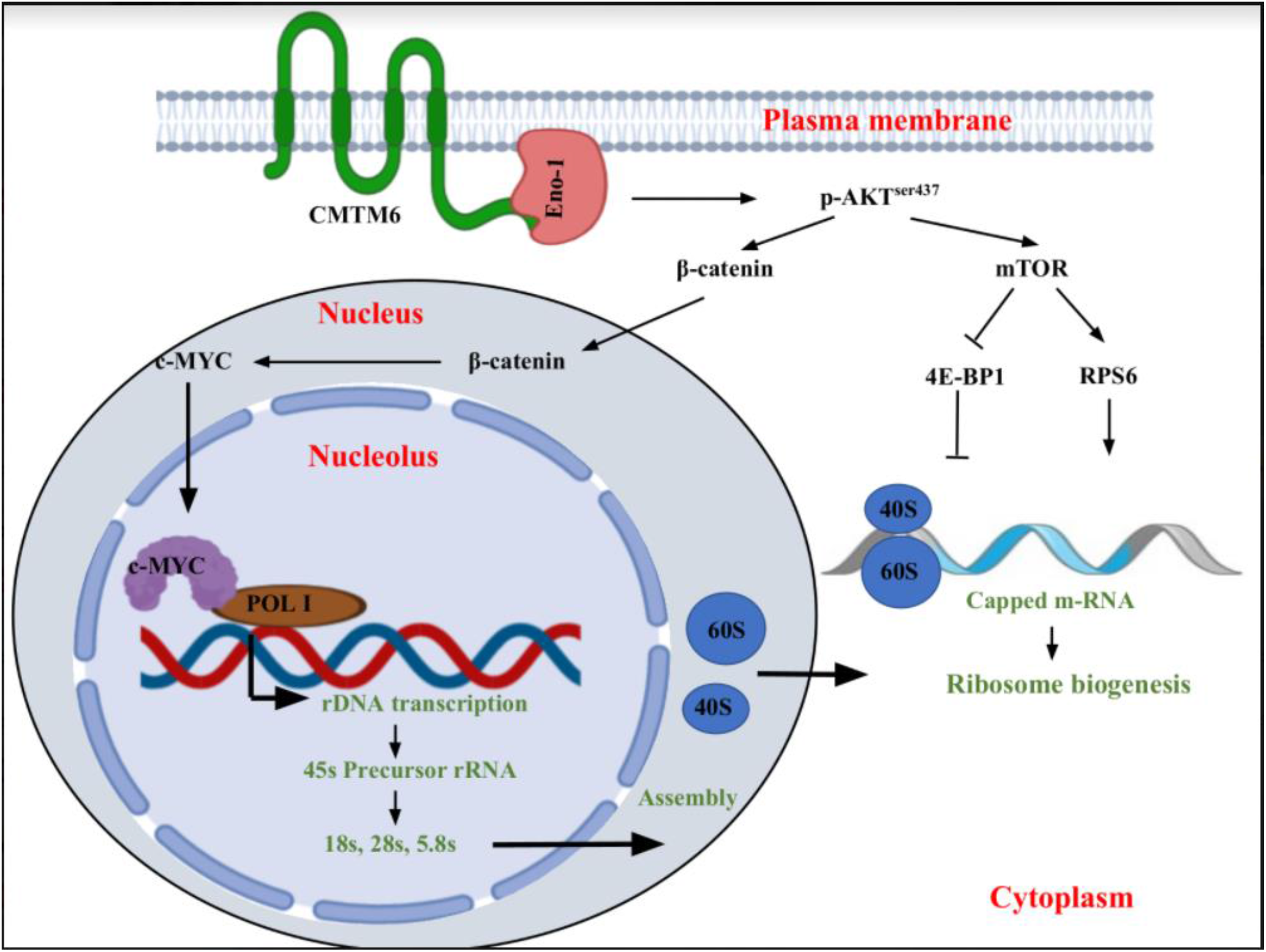

Overall, in this study, we found that CMTM6 is a novel regulator of ribosome biogenesis in chemoresistant OSCC. Blocking ribosome biogenesis through genetic inhibitor (CMTM6KD) or pharmacological inhibitor (CX-5461) can restore cisplatin mediated cell death in chemoresistant OSCC. Hence, targeting ribosome biogenesis may be viable approach to overcome chemoresistance. The novel combination of cisplatin and CX-5461 needs clinical investigation in advanced OSCC.

## Materials and methods

### Cell culture

The human tongue OSCC lines (H357 and SCC4) were obtained from Sigma Aldrich, sourced from European collection of authenticated cell culture. All OSCC cell lines were cultured and maintained in DMEM F12 supplemented with 10% FBS (ThermoFisher Scientific Cat #12500062) and penicillin–streptomycin (Pan Biotech Cat # P06-07100).

### Lentivirus production and generation of stable CMTM6 KD cell lines

pLKO.1vector (Plasmid Cat #10878) was obtained from addgene which is kindly deposited by David Root lab (28). shRNAs targeting CMTM6 were cloned into pLKO.1vector as per the protocol mentioned by addgene. Lentiviruses were produced by transfection of pLKO.1vector plasmid along with packaging plasmid psPAX2 and envelop plasmid pMD2G into HEK293T cells (29). All shRNA sequences used in this study are mentioned (Supplementary table 2).

### Transient overexpression of CMTM6, c-MYC and myr-AKT in CMTM6 KD cell lines

CMTM6 knockdown cells, stably expressing shRNA#2 targeting 3’ UTR of CMTM6 mRNA, were transiently transfected with pCMV6 CMTM6 (Myc-DDK-tagged) (Origene, Cat# RC201061) using the ViaFect transfection reagent (Promega Cat# E4982). The transfection efficiency was confirmed by immunoblotting against Anti-CMTM6 and Anti-DDK. 901 pLNCX myr HA Akt1 (Addgene, Cat #9005) was used for transient overexpression of active AKT. The myr HA Akt1 vector was kindly deposited to Addgene by Sellers WR laboratory. pCDNA3-HA-HA-humanCMYC (Addgene, Cat #74164)was used for c-MYC overexpression, which was kindly deposited by Martine Roussel Lab.

### Immunoblotting

Cell lysates were used for immunoblotting experiments as described earlier (30). For this study, we used antibody against β-actin (Sigma-Aldrich, Cat#A2066), CMTM6 (Sigma-Aldrich, Cat#HPA026980), DDK (CST: Cat#14793), AKT(CST, Cat #9272S), pAKT(Ser473) (CST, Cat #4058S), c-Myc(CST, Cat # 9402), m-TOR (CST, Cat #2983s), p-m-TOR (Ser2448) (CST, Cat #5536s), p-m-TOR (Ser2481) (CST, Cat #2974s), p-4E-BP1 (CST, Cat #2855s), RPS6 (CST, Cat #2217s), p-RPS6 (CST, Cat #5364s) and Anti-puromycin (Sigma-Aldrich, Cat #MABE343). The band intensity in all immunoblots (n=3) were quantified using imageJ software. The mean value of band intensity is indicated under the immunoblots. The intensity for each blot was calculated by normalizing with the intensity of β-actin blot.

### Cell viability assay

Cell viability was measured by 3-(4, 5-dimethylthiazol-2-yl)-2, 5-diphenyltetrazolium bromide (MTT; Sigma-Aldrich) assay as per manufacturer’s instruction.

### Colony formation assay

Colony formation assay was performed as described in Shriwas et al (31).

### Annexin-V PE/7-AAD Assay

Apoptosis and cell death assay was performed by using Annexin V Apoptosis Detection Kit PE (eBioscience™, USA, Cat # 88-8102-74) as described earlier and cell death was monitored using a flow cytometer (BD FACS Fortessa, USA).

### RT-PCR and Real Time Quantitative PCR

Direct-zol RNA Miniprep (Zymo research, Cat# R2051) was used to isolate total RNA as per manufacturer’s instruction and quantified by Nanodrop. c-DNA was synthesised by reverse transcription PCR using Verso cDNA synthesis kit (ThermoFisher Scientific, Cat # AB1453A) from 300 ng of RNA. qRT-PCR was carried out using SYBR Green master mix (Thermo Fisher scientific Cat # 4367659). GAPDH was used as a loading control.

### Immunohistochemistry

Immunohistochemistry of formalin fixed paraffin-embedded samples (OSCC patients tumor and Xenograft tumors from mice) were performed as previously describe (17). Antibodies against CMTM6 (Sigma-Aldrich, Cat#HPA026980), and Ki67 (Vector, Cat #VPRM04) were used for IHC. Images were obtained using Leica DM500 microscope. Q-score was calculated by multiplying percentage of positive cells with staining (P) and intensity of staining (I). P was determined by the percentage of positively stained cells in the section and I was determined by the intensity of the staining in the section i.e. strong (value=3), intermediate (value=2), weak (value=1) and negative (value=0).

### Immunofluorescence

The cells were seeded on lysine coated coverslip and cultured for overnight. Cells were fixed with 4% formaldehyde for 15 min, permeabilized with 1 × permeabilization buffer (eBioscience, Cat #00-8333-56) followed by blocking with 3% BSA for 1 h at room temperature. Then the cells were incubated with primary antibody overnight at 4 °C, washed three times with PBST followed by 1hr incubation with Goat anti–Rabbit IgG(H+L) secondary Antibody, Alexa Fluor® 488 conjugate (Invitrogen, Cat #A -11008). After final wash with PBST (thrice) coverslips were mounted with DAPI (Slow Fade ® GOLD Antifade, Thermo Fisher Scientific, Cat # S36938). Images were captured using a confocal microscopy (LEICA TCS-SP8). Anti CMTM6 (Sigma-Aldrich, Cat#HPA026980), Anti Fibrillarin (CST, Cat #8814) were used in this study.

### Zebrafish xenograft

The experimental protocols used for this work were approved by the institutional animal ethical review committee. PDC1 control and CMTM6 knockdown stable cells were suspended in individual tubes at a density of 1x 106cells/ml in normal media followed by addition of 5μl of the cell-labeling solution (Vybrant™ DiI Cell-Labeling Solution Catalog number: V22885) and mixed well by gentle pipetting. The cells were incubated for 20 minutes at 37°C to obtain uniform labeling and then centrifuged at 1100 rpm for 5 minutes at RT. The supernatant was removed and cells were resuspended gently in 1ml of warm(37°C) media and centrifuged at 1100 rpm for 2 minutes for the removal of extra dye and the wash step was repeated twice. The Dil stained cells were resuspended at final density 200cells/nl and ∼400 cells were microinjected (Femtojet microinjector) into perivitelline space of 48 hpf embryos of zebrafish (Danio rerio) [Tg(fli1:nEGFP)] for the development of tumor. The images of zebrafish embryos were captured using a fluorescence stereomicroscope (Leica MZ16) on the day of injection (Day 0), followed by Cisplatin treatment (500ng/ml) and CX-5461(500nM) treatment in respective groups, 3 days post injection (Day 3) and then final imaging 5 days after injection (Day 5). The tumor growth was assessed by an increase or decrease in fluorescence intensity on the 5th day compared to the day of injection (Day 0). The quantitation of fluorescence intensity was performed using ImageJ software and represented as mean fluorescent intensity where day 0 reading was taken as baseline.

### Patient Derived Xenograft

BALB/C-nude mice (6-8 weeks, male, NCr-Foxn1nu athymic) were purchased from Hylasco Bio-Technology (India) Pvt. Ltd (Telengana, India). For xenograft model early passage of patient-derived cells (PDC) established from chemo non-responder patient (treated with TPF without having any response) was considered. Two million cells were suspended in phosphate-buffered solution-Matrigel (1:1, 100 μl) and transplanted into upper flank of mice. The PDC WT cells were injected in right upper flank of the mice. These mice were randomly divided into 4 groups (n=5) once the tumors reached a volume of 50 mm3 and injected with vehicle, Cisplatin (3mg/kg), CX-5461 (10mg/kg) and Cisplatin (3mg/kg) combined with CX-5461 (10mg/kg) intraperitonially twice a week. Tumor size was measured using digital Vernier caliper twice a week until the completion of experiments. Tumor volume was determined using the following formula: Tumor volume (mm3) = (minimum diameter)2 × (maximum diameter)/2.

### OSCC patient sample

All samples collected were locoregionally advanced OSCC. Neoadjuvant CT had been prescribed before surgery and/or radiotherapy. The 3-drug combination like TPF was having highest response (TAX 324). However, some cases had poor Eastern Cooperative Oncology Group performance status; hence 2 drugs (TP) were prescribed instead of TPF. After CT the response was evaluated as per Response Evaluation Criteria In Solid Tumors criteria by clinical and radiological evaluation. If there was no evidence of malignancy, then it was diagnosed as complete response (CR). If the target lesions had decreased more than equal to 30% of the sum of the longest diameter, then it was diagnosed as partial response (PR). If there was no sign of either CR or PR, then it was called stable disease, and if the target lesions had increased more than or equal to 20% of the sum of the longest diameter, then it was called PD (progressive disease). As the patients showing CR and PR responded to the CT, they were categorized as responders, and the patients with stable disease or PD with almost no response to CT were categorized as nonresponders. The Human Ethics Committee (HEC) of the Institution of Life Sciences approved all patient-related studies, and informed consent was obtained from all patients. Study subject details with treatment modalities are presented in Supplemental Table 1.

### Puromycin labelling for protein translation determination

To determine the translation rate, cells were plated in 10 cm dish on the day before the puromycin labelling. Next day, cells should be actively growing with desired confluence of 70–80%. For labelling, 10 μM of puromycin was added directly to cell culture dish, mixed well, and placed in incubator at 37 °C for 15 min. Puromycin was washed off with PBS and the cells were collected for protein extraction and immunoblotting.

### Correlation analysis of ribosome biogenesis related genes and CMTM6

The correlation analysis was performed between ribosome biogenesis related genes and CMTM6 in HNSCC patient tumors using GEPIA (http://gepia.cancer-pku.cn/detail.php?gene=CMTM6) online analysis software based on the TCGA database and Genotype, using |log2FC|≥1 and a P value of less than or equal to 0.05 as the cut-off criteria.

### Statistical analysis

All data points are presented as mean and standard deviation and Graph Pad Prism 5.0 was used for calculation. The statistical significance was calculated by one-way variance (one-way ANOVA), Two-Way ANOVA and considered significance at P≤0.05.

