## Supplementary Figures for "CMTM6 mediates cisplatin resistance in OSCC by regulating AKT/c-MYC driven ribosome biogenesis"

### Supplementary Figure 1

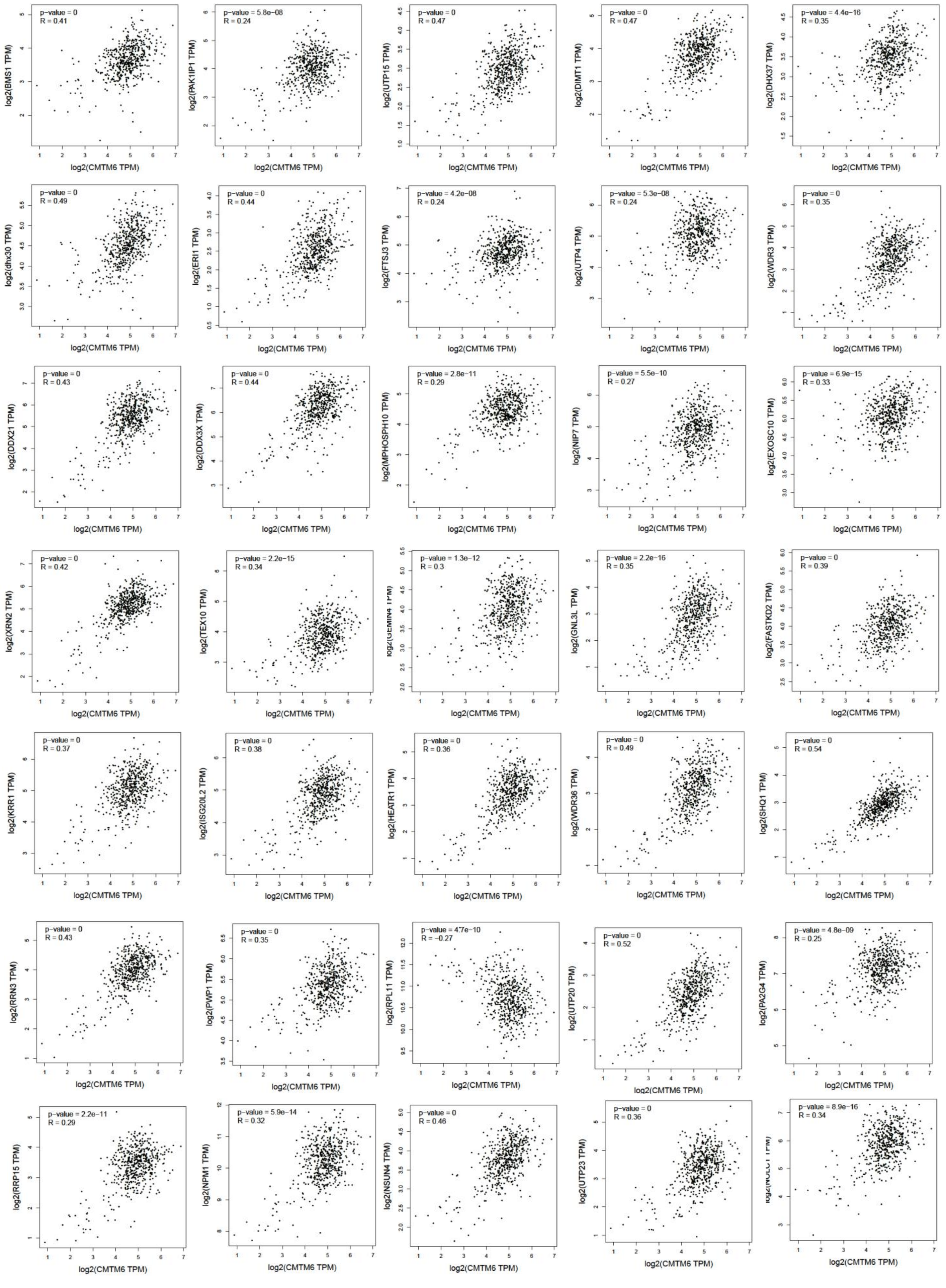

**Supplementary figure 1: CMTM6 correlates with Ribosome biogenesis related genes:** Expression correlation between CMTM6 and Ribosome biogenesis related genes RNA expression in the TCGA HNSCC database. Correlation was analyzed using Spearman's correlation coefficient test, n = 520. The analysis was performed in Gene expression profiling interactive analysis (GEPIA) platform

### Supplementary Figure 2

**A**

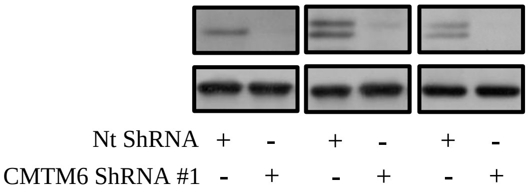


**B**

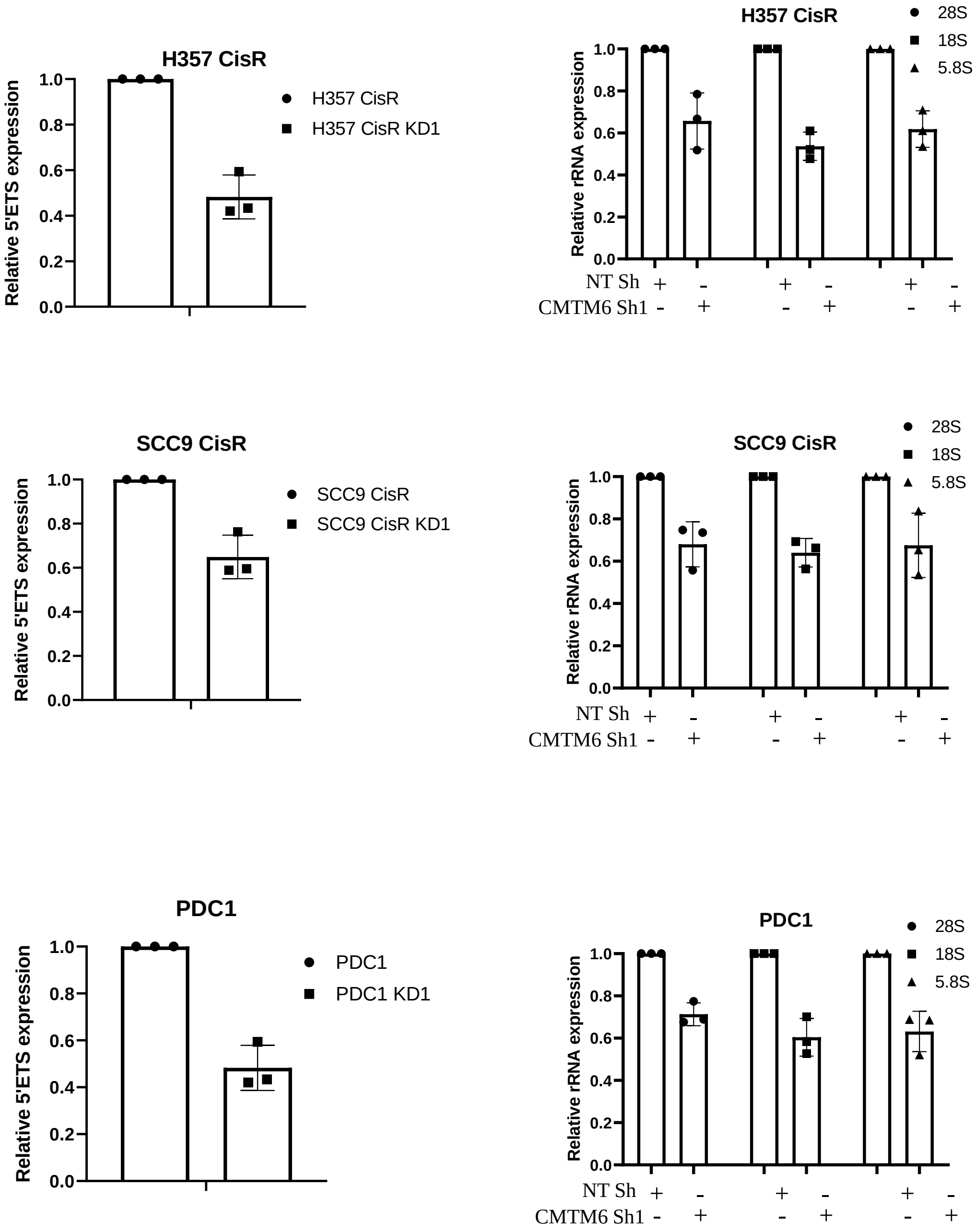


**Supplementary figure 2: CMTM6 regulates rRNA transcription:** A) Cisplatin resistant OSCC lines were stably transfected with NTShRNA and CMTM6ShRNA. ShRNA#1 targets CDS of CMTM6 mRNA. Immunoblotting (n=3) was performed with indicated antibody of indicated genes. B) Relative 5'ETS and rRNA (fold change) expression analyzed by qRT PCR in indicted cells (mean ±SEM, n=3) , *: $P < 0.05$ by two way ANOVA.

**Supplementary Figure 3**

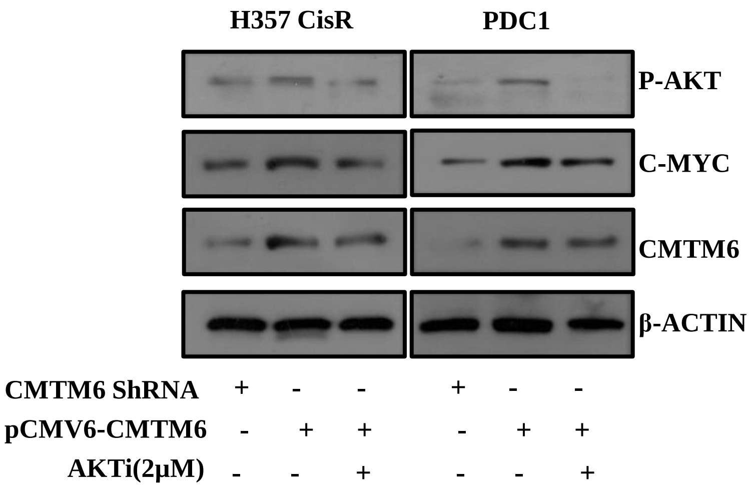


**Supplementary figure 3:** CMTM6 was overexpressed in chemoresistant cells stably expressing CMTM6ShRNA#2 and AKTi was treated. Immunoblotting was performed with indicated antibodies

**Supplementary Figure 4**

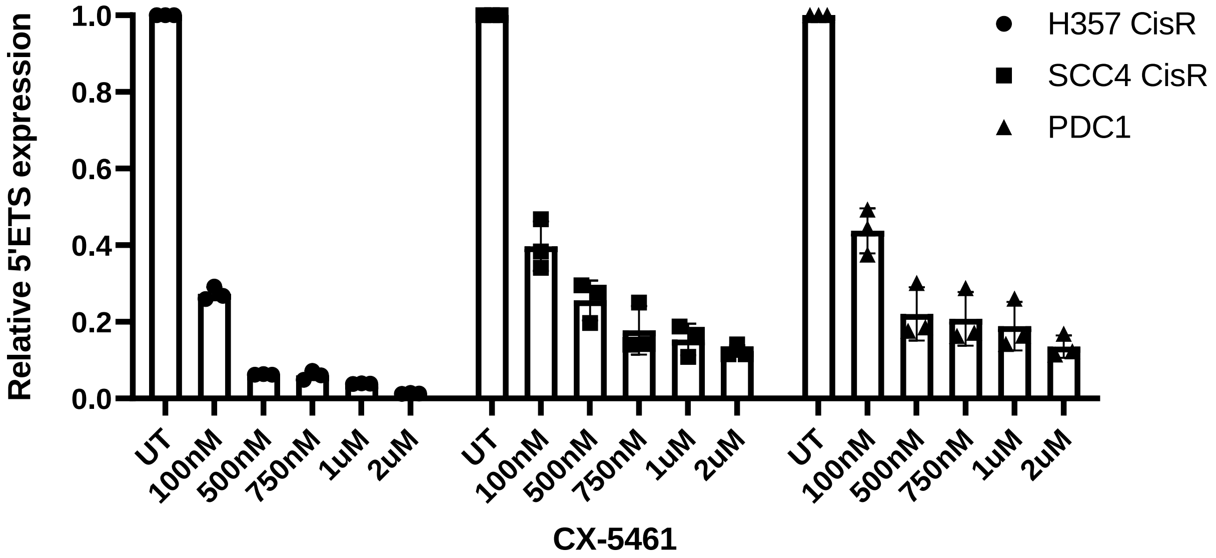


**Supplementary figure 4:** Indicated cells were treated with CX-5461 for 48h with indicated concentration and relative 5'ETS expression analyzed by qRT PCR (mean ±SEM, n=3) , *: $P < 0.05$ by two way ANOVA.

**Supplementary Figure 5**

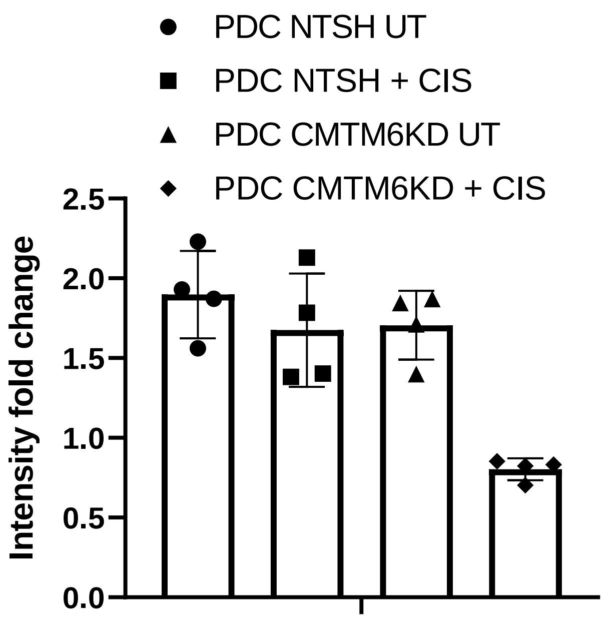


**Supplementary figure 5:** The tumor growth was assessed by an increase in fluorescence intensity on the 5th day compared to the day of injection. n=4 and the quantitation of fluorescence intensity was performed using ImageJ software.
